# Birth of protein-coding exons by ancient domestication of LINE retrotransposon

**DOI:** 10.1101/2024.04.25.591049

**Authors:** Koichi Kitao, Kenji Ichiyanagi, So Nakagawa

**Affiliations:** Laboratory of Genome and Epigenome Dynamics, Department of Animal Sciences, Graduate school of Bioagricultural Sciences, Nagoya University, Furo-cho, Chikusa-ku, Nagoya 464-8601, Japan; Depertment of Molecular Life Science, Tokai University School of Medicine, Isehara, Kanagawa 259-1193, Japan; Division of Genome Sciences, Institute of Medical Sciences, Tokai University, Isehara, Kanagawa 259-1193, Japan; Division of Interdisciplinary Merging of Health Research, Micro/Nano Technology Center, Tokai University, Hiratsuka, Kanagawa 259-1292, Japan

**Author notes:** Koichi Kitao; So Nakagawa, (KK); (SN).

**Keywords:** *de novo* gene birth, LINE, alternative splicing, retrotransposon, Sauria

## Abstract

Transposons, occasionally domesticated as novel host protein-coding genes, are responsible for the lineage-specific functions in vertebrates. LINE-1 (L1) is one of the most active transposons in the vertebrate genomes. Despite its abundance, the contribution of L1 to the birth of vertebrate proteins remains unelucidated. Here, we present a novel mechanism for the origination of *de novo* proteins, in which the L1 retrotransposons are incorporated into host genes as protein-coding exons by alternative splicing. L1 ORF1 protein (ORF1p) is an RNA-binding protein that binds to L1 RNA and is required for retrotransposition by acting as an RNA chaperone. We identified a splicing variant of *myosin light chain 4* (*MYL4*) containing an L1 ORF1-derived exon and encoding a chimeric protein of L1 ORF1p and MYL4, named Lyosin. Molecular evolutionary analysis revealed that Lyosin was acquired in the common ancestor of reptiles and birds during the Paleozoic era. The amino acid sequence of Lyosin had undergone purifying selection although it was lost in some lineages, including the Neognathae birds and snakes. The transcripts encoding Lyosin were expressed in the testes of two lizard species, suggesting that its function is different from that of the canonical MYL4 expressed specifically in the heart. Furthermore, sequence searches revealed other evolutionarily conserved chimeric isoforms fused to the L1 ORF1p in three genes in vertebrates. Our findings suggest a novel evolutionary mechanism for the birth of lineage-specific proteins derived from transposons and implicate the previously unrecognized adaptive functions of L1 ORF1p.

## Introduction

The emergence of lineage-specific proteins is important for the diversified functions of vertebrates. Vertebrate genomes are occupied by a large number of transposons, accounting for several percent to tens of percent of host genomes (1). Occasionally, transposon-encoded proteins have been “domesticated” as novel host proteins (2). For example, Syncytin proteins encoded by *env* genes of endogenous retroviruses (ERVs) contribute to trophoblast cell fusion and immunosuppression in the mammalian placenta (3, 4). The *gag* genes encoding virus-like capsid proteins of long terminal repeat (LTR) retrotransposons are also co-opted as host genes: *ARC* for synaptic plasticity (5, 6), *Peg10* and *Rtl1* for placenta formation (7, 8), and *PNMA2* expressed in neurons and related with autoimmune diseases (9). Other examples include *ASPRV1/SASPase*, derived from a retroviral protease involved in the protease activity for mammalian skin formation (10); *NYNRIN*, derived from integrase and contributing to placentation (11); and *RTOM* family, derived from reverse transcriptase in monotremes (12). DNA transposons are also sources of multiple lineage-specific proteins. The co-option events of DNA transposons as exons of host transcription factors have occurred recurrently throughout vertebrate evolution and are thought to facilitate the evolution of species-specific transcriptional networks (13).

Long interspersed nuclear elements (LINEs) are non-LTR retrotransposons that are found in various vertebrates (1). LINE-1 (L1) is the only active LINE in the human genome and has two open reading frames (ORFs), ORF1 and ORF2. ORF1 encodes the ORF1p, an RNA-binding protein required for retrotransposition. ORF2p, encoded by ORF2, is an enzymatic protein that contains the endonuclease and reverse transcriptase domains. Some LINE clades do not have non-enzymatic proteins unlike L1 ORF1p (14). Despite their abundance in vertebrate genomes, few examples of LINE domestication for novel protein-coding genes have been reported. *L1TD1* consists of two coding exons derived from L1 ORF1 and is conserved in eutherian mammals. This gene was first identified in a screen for reprogramming-related proteins in mouse ES cells, but knockout of *L1td1* showed no phenotypic effects on cell reprogramming or mouse development (15). In contrast, other studies suggest that human L1TD1 is involved in cell reprograming (16). Mechanistically, L1TD1 is involved in post-transcriptional regulation by controlling intracellular protein condensation (17). The other example is a ruminant-specific gene, *BCNTp97*, derived from the tandem duplication of *BCNT* gene followed by insertion of a retrotransposable element-1 (RTE-1) (18). RTE-1 is classified as LINE superfamily (14). The *BCNTp97* encodes an intact endonuclease domain derived from RTE-1; however, its molecular function remains unknown (18). Although these few LINE-derived genes have been identified in mammals, their evolution and diversity are poorly understood.

In the present study, we identified a novel LINE-derived protein, which is encoded by a non-canonical splicing variant of *myosin light chain 4* (*MYL4*). This protein is a chimeric fusion protein of L1 ORF1p and MYL4. The expression of the chimeric isoform is tightly controlled by alternative splicing to be expressed in the testis, while maintaining the normal expression of the canonical *MYL4* isoform in the heart. We also identified three other examples of L1 ORF1p chimeric isoforms in vertebrates. This study proposes a new evolutionary model for the emergence of *de novo* genes and a previously unrecognized functional benefit of LINEs in host adaptation.

## Results

### Identification of Lyosin: an L1 ORF1p and MYL4 chimeric isoform

According to RefSeq annotation, American alligator (*Alligator mississippiensis*) *MYL4* has a non-canonical exon (hereafter called “exon L”) that is not present in mouse (**Fig. 1A**). We found that the protein encoded by the exon L is similar to L1 ORF1p (**Fig. 1B**). The amino acids encoded by exon L show 30% identity and 51% similarity to L1 ORF1p. Exon L was predicted to be connected to exon 3 instead of the canonical exon 1 and 2. Therefore, this splicing variant was inferred to encode a chimeric protein of L1 ORF1p and MYL4. We named this putative protein Lyosin (L1-myosin light chain 4 chimeric protein). The L1 ORF1p consists of four domains: the disordered N-terminal domain (NTD), coiled-coil (CC), RNA recognition motif (RRM), and the C-terminal domain (CTD) (19). This domain arrangement is classified as type II ORF1p among LINE families and is not identical to any other RNA-binding proteins (20). Structural prediction revealed that Lyosin retained domains corresponding to the NTD, CC, and RRM (**Fig. 1C**). The predicted RRM structure of Lyosin showed a characteristic βαβββαβ structure observed the L1 ORF1p RRM domain (**Fig. 1D**). Moreover, superposition of the predicted Lyosin RRM domain with the L1 ORF1p RRM domain determined by X-ray diffraction of the crystal structure (20) showed a high degree of similarity (RMSD = 2.914Å) (**Fig. 1E**). These structural analyses suggest that the protein encoded by exon L is homologous to L1 ORF1p. MYL4 and other myosin light chains belong to the EF-hand calcium-binding protein family (21) although MYL4 itself has lost its calcium-binding capacity (22). The protein sequence encoded by exon 3 and the later exons included in Lyosin corresponds to the EF-hand calcium-binding motifs (**Fig. 1B**). Taken together, these structural analyses indicate that Lyosin is a chimeric protein composed of the NTD, CC, RRM domains of L1 ORF1p and the EF-hand domain of MYL4.

**Fig. 1.**
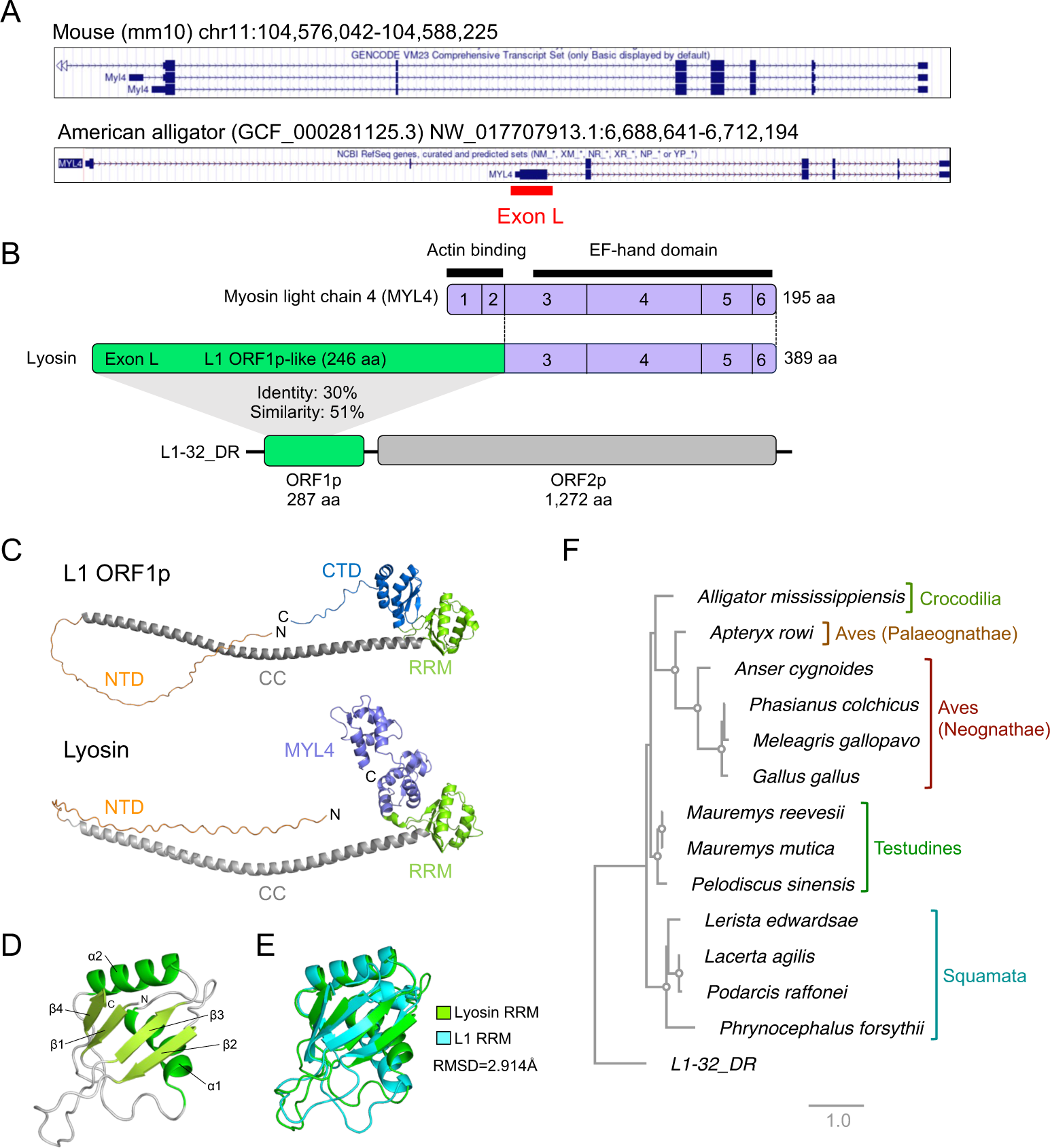
Lyosin in reptiles and birds. (A) UCSC genome browser of *MYL4* of mouse and American alligator. The non-canonical exon L was annotated in American alligator’s *MYL4*. (B) Representation of the MYL4 and Lyosin gene structure in American alligator. The amino acid sequences of exon 1 to 2 and exon 3 to 6 correspond to the actin-binding site and the EF-hand calcium-binding domain, respectively, in MYL4. Lyosin consists of exon L and exon 3 to 6 of MYL4. Exon L encodes a 246 aa protein similar to L1 ORF1 protein (ORF1p), which is an RNA-binding protein. The ORF2 protein (ORF2p) of L1 contains an enzymatic protein with endonuclease and reverse transcriptase activity. (C) Protein structures of human L1 ORF1p (PDB: AF_AFQ9UN81F1) and alligator Lyosin predicted by AlphaFold2. ORF1p consists of the disordered N-terminal domain (NTD), coiled-coil (CC), RNA recognition motif (RRM), and C-terminal domain (CTD). Lyosin contains NTD, CC, and partial RRM. (D) The βαβββαβ structure of the predicted Lyosin RRM domain. (E) Structural comparison of the predicted Lyosin RRM domain and the L1 ORF1p RRM domain determined by X-ray diffraction (PDB: 2W7A). (F) Maximum likelihood-based phylogenetic tree of Lyosin proteins obtained by NCBI BLASTP search. Open circles in internal nodes indicate >95% ultrafast bootstrap support (1,000 replicates).

### Identification of Lyosin in reptiles and birds

Next, we investigated whether Lyosin is present in any species other than the American alligator. We performed a BLASTP search against the NCBI non-redundant (nr) protein database using the amino acid sequence of the alligator Lyosin as a query. The search identified Lyosin-like proteins in three species of Testudines (turtles), four species of Squamata (lizards and snakes), and five species of Aves (**Dataset S1**). Alignment of these proteins revealed that the sequences corresponding to L1 ORF1p were divergent, whereas those of MYL4 were highly conserved. In particular, deletions in the region corresponding to the RRM domain of the MYL4 region were observed in birds, except Okarito kiwi (*Apteryx rowi*) (**Fig. S1**). We then constructed a molecular phylogenetic tree of L1 ORF1p and Lyosin-like protein exon L to gain insights into their evolutionary relationships. The maximum-likelihood based tree showed that the exon L of Lyosin-like proteins was distinct from L1 ORF1p and further reflected the host evolution (**Fig. 1F**). This suggests that the Lyosin was captured and evolutionarily conserved from their common ancestor.

### *Lyosin* in tetrapoda genomes

Lyosin is encoded by a non-canonical splicing variant, and it is probable that Lyosin has not been annotated in most genomes, resulting in the absence of protein databases. Therefore, we conducted a comprehensive genome-wide search for *Lyosin* sequences. First, we performed an intron-considered BLAT search on tetrapod genomes using the full-length amino acid sequences of Lyosin as queries (**Fig. 2A**). Approximately 40% of the Lyosin corresponds to canonical MYL4, and approximately 60% of it corresponds to L1 ORF1p-like amino acids. Therefore, we set the threshold for Lyosin detection to a query coverage of 50% or more. As a result of this search, *Lyosin* sequences were detected in the genomes of Aves (185 out of 556 genomes), Crocodilia (4 out of 4 genomes), Testudines (25 out of 25 genomes), Sphenodon (1 out of 1 genome), and Squamata (37 out of 49 genomes), but not in those of mammals and amphibians (**Fig. 2A; Dataset S2**). This result is consistent with the previous results of the BLASTP search for the NCBI nr protein database. This confirms that *Lyosin* was acquired in the ancestor of Sauria, the crown group of birds and reptiles, at least before 280 million years ago, during the Paleozoic era. *Lyosin* was not detected in several species of Aves and Squamata. It is also possible that coding capacities or splicing sites of the *Lyosin* exon L were lost even in species where *Lyosin* sequences were detected by BLAT search. Therefore, we considered those that retained canonical 5’ splice sites (5’-GT-3’) and encoded more than 200 aa amino acids as the intact exon L. Among the genomes in which *Lyosin* was detected by BLAT search, the intact exon L was present in the genomes of Aves (16 out of 185 genomes), Crocodilia (4 out of 4 genomes), Testudines (23 out of 25 genomes), and Squamata (14 out of 37 genomes) (**Fig 2A**). Crocodilia was the only group in which all species retained the intact exon L. These genomic searches suggest that *Lyosin* was repeatedly lost during the evolution of Sauria.

**Fig. 2.**
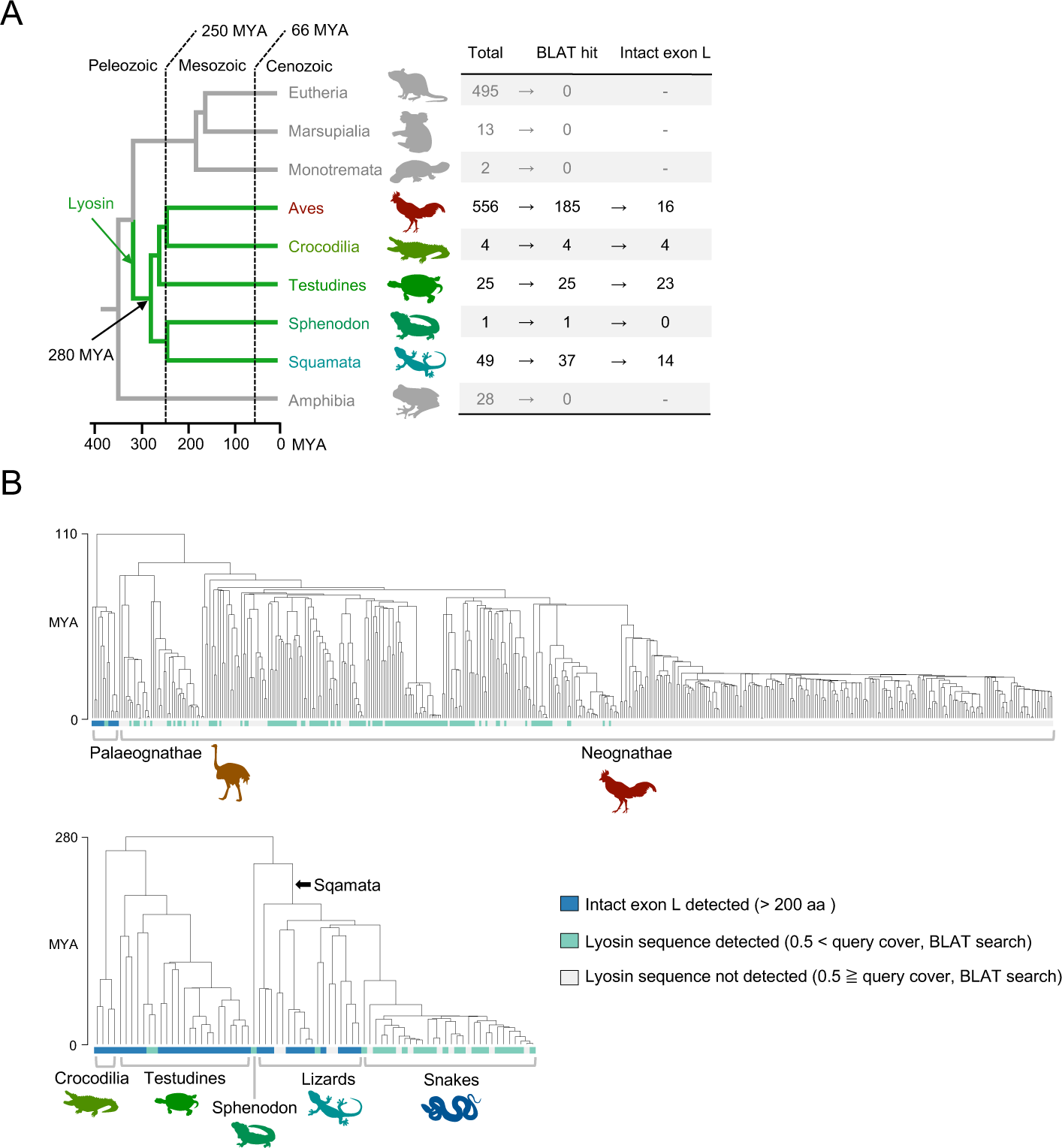
Identification of *Lyosin* in tetrapoda genomes. (A) Schematic summary of the detection of *Lyosin* in tetrapoda genomes. The numbers of analyzed genomes (Total), *Lyosin*-detected genomes by BLAT searching (BLAT hit), and intact *Lyosin*’s exon L-detected genomes are listed on the right side. All *Lyosin*-detected species were included in the Sauria clade, indicating that the *Lyosin* co-option was occurred in its common ancestor before 280 million years ago (MYA) in the Paleozoic era. (B) The evolutionary history of *Lyosin* in reptile and bird lineages. Phylogenetic trees of species with divergence times were based on the TimeTree (Kumar et al. 2022). The results of *Lyosin* detection (**Dataset S2**) were reflected on the tree nodes.

### Multiple losses of *Lyosin* in the Sauria clade

To track the losses of *Lyosin*, we mapped the presence or absence of *Lyosin* onto a species tree obtained from the TimeTree database (23). First, Aves was analyzed separately from the other reptilian clades because of the large number of species. Tracing *Lyosin* in Aves, we found that the intact exon L of *Lyosin* was retained only in the Palaeognathae species, and none of the Neognathae species had the intact exon L in (**Fig. 2B**). This is consistent with the observation that the most RefSeq proteins of the Neognathae species contain deletions in the exon L region of *Lyosin* (**Fig. S1**), suggesting that *Lyosin* has not been under evolutionary conservative selection in Neognathae. In reptiles, losses of *Lyosin* occurred at least once in Testudines and Sphenodon, independently, and three times in Squamata (**Fig. 2B**). Notably, no Serpentes (snakes) in Squamata retained the intact exon L. Although some analyzed species were not shown in the tree of Fig. 2B due to the lack of those species in the TimeTree database (23), we confirmed that neither Neognathae nor Serpentes species, including the ones missed in the tree, contain intact exon L (**Dataset S2**), Taken together, losses of *Lyosin* have occurred independently during the Sauria evolution.

### Molecular evolution of amino acid sequences in Lyosin

Frequent losses of *Lyosin* raises the suspicion that the exon L of *Lyosin* has remained by chance and is not evolutionarily conserved as a functional protein-coding exon. Therefore, we next investigated the conservation of the amino acid sequences of exon L. Alignment of the amino acid sequences revealed that the conservation level differed in protein domains. In lizards, where the sequences were relatively diverse, the amino acids were conserved at the N-terminus of the CC domain and at both termini of the RRM domain (**Fig. 3**). The same trend was also observed for Palaeognathae and Testudines. As for the RRM domain, the conservation levels were high at the structured α-helix and β-sheet, whereas those were relatively low at the loop region (**Fig. S3**). These data suggest that the functional constraints of the Lyosin protein structure dictated their amino acid sequence conservation. We next conducted molecular evolutionary analyses on the amino acid sequence of exon L.

**Fig. 3.**
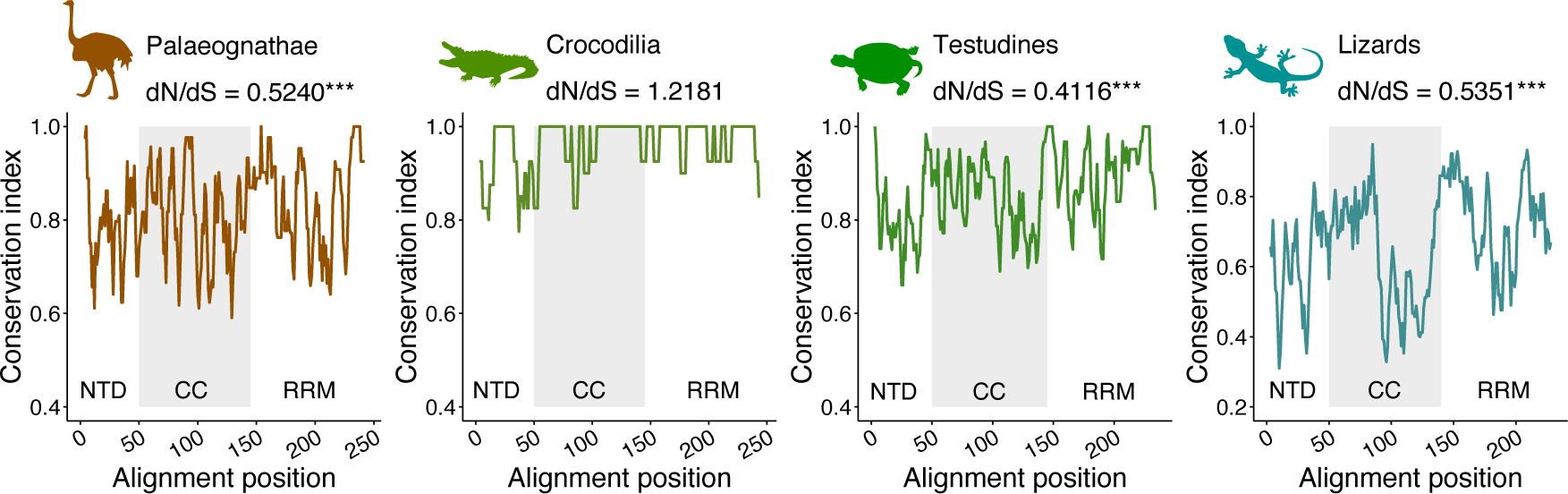
Evolutionary conservation of each amino acid site in the Lyosin protein. Amino acid sequences encoded in exon L were aligned. The conservation index was calculated by summing of the squares of the amino acid rates at each site. Sites with gaps were excluded. Plots were obtained along a sliding window of five sites for the conservation index. The dN/dS values were analyzed by the likelihood ratio test; ***p < 0.001.

We calculated the ratio of nonsynonymous and synonymous codon substitution (dN/dS) in species that retained the intact exon L (**Dataset S2**). The dN/dS ratio was less than one, suggesting a purifying selection of amino acids in Palaeognathae, Testudines, and lizards (**Fig. 3**). However, in Crocodilia, the dN/dS ratio was found to be greater than 1. This may be due to the relatively shallow branching in Crocodilia species and the very slow rate of base substitution in Crocodilia (24), resulting in insufficient nucleotide substitutions to examine selective pressure. We also evaluated positively selected amino acid sites in the exon L. However, none of the sites were statistically supported by two different evolutionary models based on the likelihood ratio test, making the positive selection hypothesis less reliable (**Fig. S2**). Together, these site-specific conservation and purifying selections on the amino acids suggest that exon L of the Lyosin protein was maintained as a protein-coding exon.

### Testis-specific expression of *Lyosin* in lizards

To obtain insights into the physiological function of Lyosin, we investigated its tissue expression. *MYL4* is known to be expressed specifically in the heart of adult mammals (22). We analyzed the transcriptome data set of green anole (*Anolis carolinensis*), which retains the intact exon L of *Lyosin*. Similar to mammals, *MYL4* was highly expressed in the green anole heart, whereas low expression was observed in several other tissues (**Fig. 4A**). To examine the expression level of the *Lyosin* splicing variant, we counted the RNA-seq reads spanning the introns. While canonical splicing between exon 2 and exon 3 was detected dominantly in heart (**Fig. 4B**), splicing between exon L and exon 3 for *Lyosin* was specifically identified in the testis (**Fig. 4C**). The genome browser view of the RNA-seq reads mapped to the *MYL4* locus also confirmed the selective expression of *Lyosin* in testis (**Fig. 4D and E**). To determine whether this expression pattern is similar in other species, reverse transcription PCR (RT-PCR) was performed on the heart and testis of Madagascar ground gecko (*Paroedura picta*) (25), which is also predicted to retain the intact exon L of *Lyosin*. The canonical *MYL4* transcripts were detected in the heart, whereas the *Lyosin* transcripts were detected in the testis (**Fig. S4A**).

**Fig. 4.**
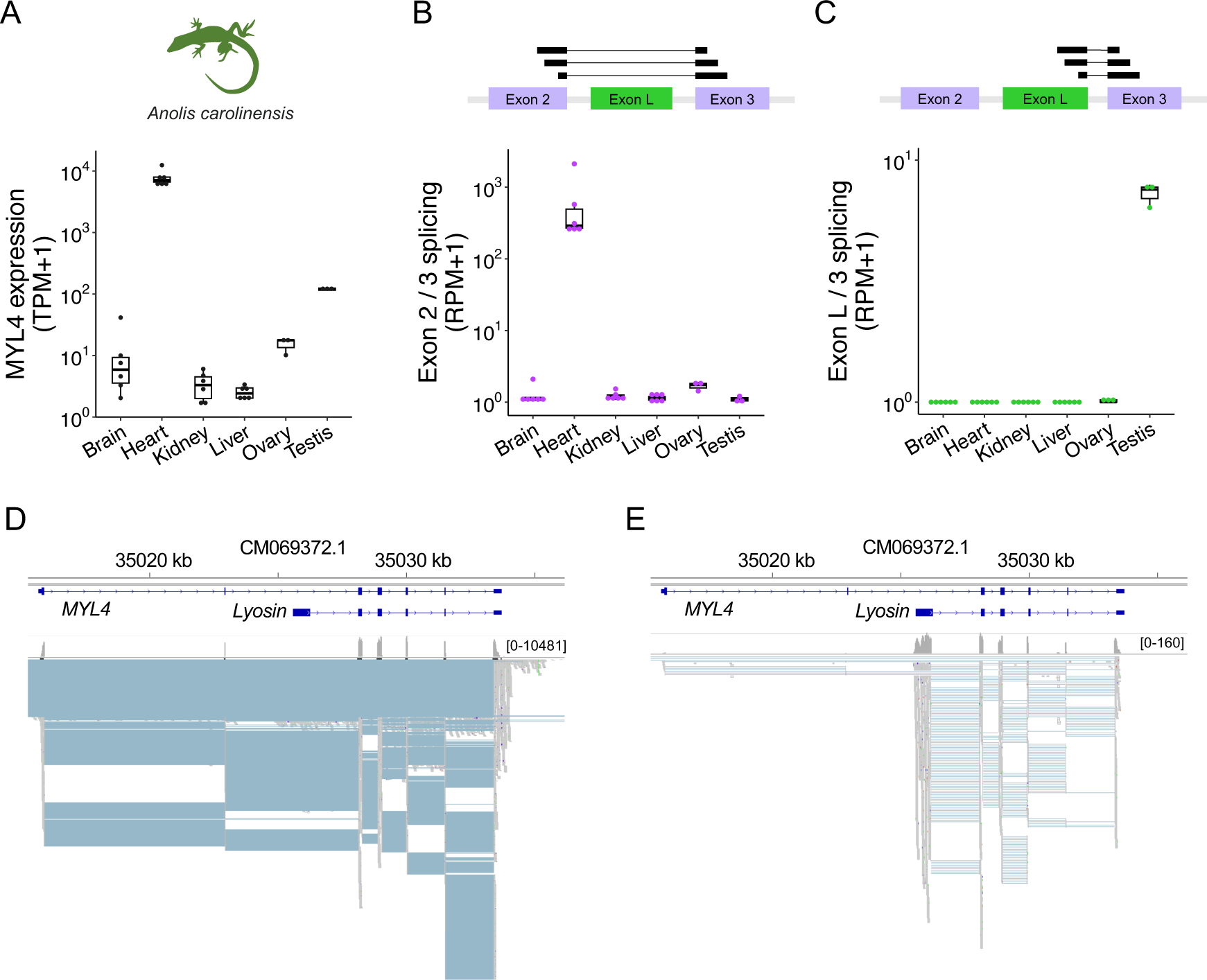
Tissue expression of *Lyosin*. (A) Box plot and point representations of the transcripts per million (TPM) of *MYL4* expression obtained from RNA-seq data of the green anole tissues. (B and C) Box plot and point representations of splice junction reads spanning exons. The read counts were normalized as reads per million (RPM): reads spanning exon 2 and 3 (B) and reads spanning exon L and 3 (C). (D and E) Genome browser view of the *MYL4* locus with RNA-seq reads of green anole. Transcripts of canonical *MYL4* and *Lyosin* were constructed by genome-guided *de novo* assembly in this study: heart (D) and testis (E).

### Molecular characterization of the Lyosin protein

We next attempted the molecular characterization of the Lyosin protein. The coding sequences for MYL4 and Lyosin of Madagascar ground gecko were cloned into a mammalian expression plasmid with the C-terminal HA tag. Then, the plasmids were transfected into the human embryonic kidney 293T cells. Western blotting analysis showed that the bands of expressed proteins were detected at the expected positions, confirming that the proteins were not cleaved (**Fig S4B**). We examined their subcellular localization in comparison with human L1 ORF1p, which is known to form the puncta required for retrotransposition (26). Unlike L1 ORF1p-HA, the Lyosin-HA was dispersed in the cytoplasm and did not exhibit the punctum formation (**Fig. S4C**). Furthermore, neither MYL4-HA nor Lyosin-HA was present in the puncta formed by co-expressed L1 ORF1p-FLAG (**Fig. S4C**). Finally, to examine the effect of the Lyosin protein on L1 retrotransposition, we performed a reporter-based L1 retrotransposition assay in 293T cells (**Fig. S4D**). The results showed that Lyosin did not affect L1 retrotransposition, whereas human MOV10 helicase, which is known to be an inhibitor of L1 mobility (27), reduced L1 retrotranspositions (**Fig. S4E**). Although it should be noted that these molecular characterizations were performed in human cells using human L1, we obtained no evidence suggesting that the Lyosin protein affects L1 activity.

### Other examples of the origination of coding exons by the ORF1p co-option

The emergence of *Lyosin* in the Sauria clade raises the possibility that the vertebrate genomes may harbor other conserved chimeric isoforms of LINE ORF1p and host proteins. To assess this possibility, we screened the vertebrate RefSeq proteins for protein-coding genes harboring both ORF1p-like and non-ORF1p-like isoforms (**Fig. 5A**) (see **Materials and Methods**). We identified four protein-coding genes with LINE chimeric isoforms, including *MYL4*. The identified ORF1p-like exons were classified as either the first or last coding exons (**Fig. S5A**). *RFX5,* which encodes DNA binding protein RFX5, had an alternative non-canonical first coding exon similar to L1 ORF1p in catfish (order Siluriformes) (**Fig. 5B**). *NUP42,* encoding nucleoporin 42, and *USP4,* encoding ubiquitin specific peptidase 4, had ORF1p-like final coding exons in Afrotheria and Carnivora, respectively (**Fig. 5C**). Structural prediction of the proteins encoded by these ORF1p-like exons confirmed the presence of CC and RRM domains as in L1 ORF1p in all three genes, and an almost complete CTD structure was confirmed in NUP42 (**Fig. S5B**). These data suggest that the emergence of novel splicing isoforms by the insertion of the L1 ORF1 and their subsequent domestication has occurred multiple times in vertebrate evolution.

**Fig. 5.**
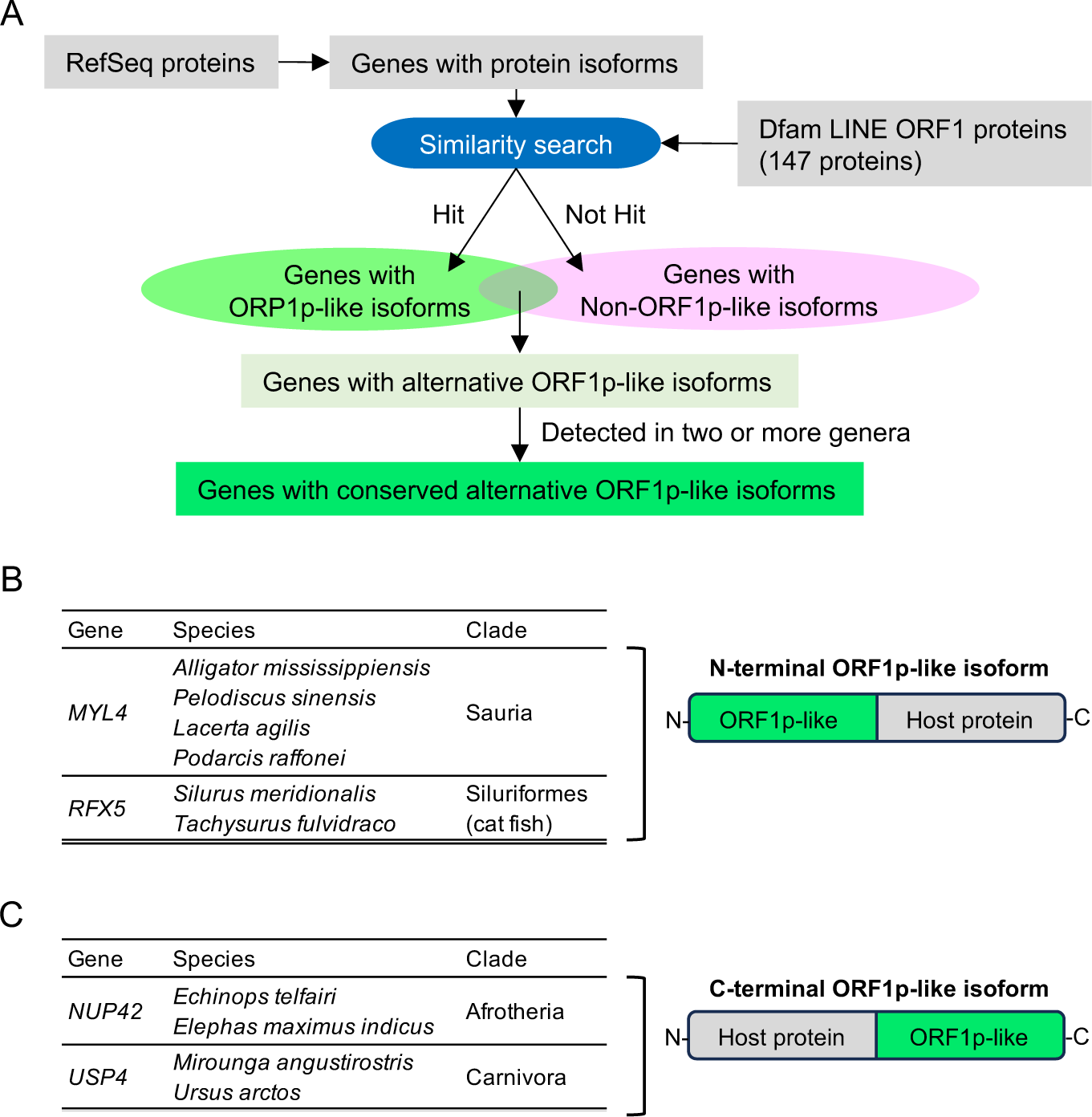
Other ORF1p chimeric isoforms in vertebrates. (A) Schematic workflow of the detection of ORF1p chimeric isoforms from RefSeq proteins. (B) In the non-mammalian vertebrate RefSeq proteins, two protein-coding genes had alternative ORF1p-like exons. In both genes, the amino acid sequences encoded by the first coding exons were similar to ORF1p. (C) In the mammalian vertebrate RefSeq proteins, two protein-coding genes had alternative ORF1p-like exons. The amino acid sequences encoded by the final coding exons were similar to ORF1p.

## Discussion

LINEs are some of the most active transposons in vertebrates. Among them, L1 is the only LINE that remains active in the human genome (28). L1 ORF1p is an RNA-binding protein that assembles to package L1 RNA (29). ORF1p then facilitates the rearrangement of nucleic acid structures and is thought to function as a nucleic acid chaperone (30). Structural analysis has suggested that both RRM and CTD are required for L1 ORF1p to bind to RNA (20). Lyosin, however, lacked CTD (**Fig. 1B**). Thus, Lyosin probably lost its native RNA-binding ability. In contrast, amino acids for the RRM domain were selectively conserved in Lyosin (**Fig. S3**), suggesting that the structure of the RRM domain itself is essential for Lyosin function. Homo-trimer formation is also important for L1 ORF1p function (31). For trimer formation in L1 ORF1p, the C-terminal half of the CC domain is necessary and sufficient (20). In Lyosin, the C-terminal half of the CC domain is less conserved than the N-terminal half, suggesting that the Lyosin could not assemble as a trimer (**Fig. 3**). Although more experimental studies are required to rationalize the molecular function and structural features of Lyosin, the discovery of Lyosin in this study suggests that proteins derived from L1 ORF1p have unrecognized biological functions other than L1 retrotransposition and will expand our understanding of the molecular functions of L1 ORF1p.

We revealed that the coding capacity of *Lyosin* has been decayed independently in several lineages (**Fig. 2**).Notably, the same recurrent gene losses have been observed in L1TD1, a mammalian L1 ORF1-derived gene. L1TD1 was acquired in the common ancestor of eutherians but was lost independently in the lineages of Afrotheria, ruminants, and bats (32). To explain this evolutionary scenario, it could be hypothesized that L1TD1 is a LINE suppressor; when the target LINE is extinct, L1TD1 becomes unnecessary and is subsequently lost by random mutations. In fact, loss of *L1TD1* has been observed in megabat lineages, in which active L1 elements are extinct (32). This hypothesis also seems to fit with the observation of the specific expression of L1TD1 in ES cells, where transposon activity should be repressed to avoid mutagenesis during early development. This also applies to the specific expression of *Lyosin* in the testis, where there is a concern that transposon insertion damages germline genomic DNA (**Fig. 4**). Nonetheless, a recent reporter-based assay showed that human L1TD1 is not capable of suppressing L1 activity (17). Similarly, there was no evidence that Lyosin represses L1 activity (**Fig. S4**). However, these analyses were validated by artificially expressed L1 from the reporter plasmids. It is also possible that L1TD1 has the potential to function as both an L1 repressor and a translational regulator, and that the balance of these dual functions differs between lineages. Lyosin provides an alternative opportunity to validate the evolutionary relationship between the frequent gene losses and the arm race against L1 elements by investigating whether the Lyosin interacts with L1 proteins and represses the L1 retrotransposition in reptilian or avian experimental systems.

Alternatively, the multiple gene losses can be explained by the gene replacement hypothesis, in which the functional replacement of one gene with another makes the existing gene indispensable and leads to its loss. Importantly, such evolutionary replacement has been proposed for the evolution of *syncytin* genes, which are derived from ERVs, another group of retrotransposons. Functional *syncytin* genes have different viral origins in mammalian lineages (33), and functional reduction of several *syncytin* genes has been reported in different mammals (34, 35). This can be explained by the scenario in which a new Syncytin inherits placental functions from older Syncytin like “baton-pass” (36). In the case of *Lyosin*, the L1 ORF1 is interspersed in the vertebrate genomes, and functional replacement of Lyosin with a new Lyosin-like protein may occur. Our analysis identified at least three ORF1p chimeric isoforms other than Lyosin (**Fig. 5**). Considering that we analyzed only the proteins included in the current RefSeq annotation, it is possible that most cases of ORF1p chimeric protein acquisition remain to be elucidated. Further detection of such protein isoforms will provide a more accurate explanation of the evolutionary mechanisms of the ORF1p-derived protein isoforms.

Genome sequencing of many vertebrates is still in progress. The identification of lineage-specific splicing isoforms remains challenging owing to the lack of orthologous relationships with model organisms. Gene prediction based on experimental data such as RNA-seq and evolutionary conservation is expected to lead to the discovery of lineage-specific splicing isoforms in the future. The discovery of Lyosin demonstrates a new mechanism of isoform evolution through the capture of a coding exon by LINE domestication. In addition, LINE insertion can alter splicing patterns by providing intrinsic transcription start sites and splicing sites (37, 38). The contribution of LINEs to the generation of lineage-specific splicing isoforms is now an open question in the era of genomic biodiversity.

## Materials and Methods

### Protein structure analysis

The protein structure of the American alligator Lyosin (XP_019356465.1) was predicted using AlphaFold2 implemented in ClabFold v1.5.2 (39). The obtained protein data bank (PDB) files of the whole Lyosin and the RRM (**Dataset S3**) were analyzed and visualized using PyMOL v2.5.0 (40).

### BLASTP search in the NCBI database

The amino acid sequence of American alligator Lyosin (XP_019356465.1) was used as a query. The BLASTP search was performed against the database of “nr All non-redundant GenBank CDS translations+PDB+SwissProt+PIR+PRF excluding environmental samples from WGS projects” with default parameters on the NCBI web server as of November 29, 2023 (**Dataset S1**). To construct the phylogenetic trees of the putative Lyosin proteins, these protein sequences were aligned using MAFFT v7.487 with the “L-INS-i” option (41). (**Dataset S4**). IQ-TREE2 v2.0.8 (Minh et al. 2020) was used to construct a phylogenetic tree with 1,000 replicates generated by an ultrafast bootstrap approximation (42).

### Tetrapoda genomic search

Tetrapoda genomes were downloaded from the NCBI assembly via GenomeSync (https://genomesync.org/) (43) accessed on January 12, 2022 (**Dataset S5**). The representative Lyosin amino acid sequences from the RefSeq database were used as queries as follows: XP_019356465.1 (*Alligator mississippiensis*), XP_025041760.1 (*Pelodiscus sinensis*), XP_025915001.1 (*Apteryx rowi*), and XP_033026289.1 (*Lacerta agilis*) for the search using BLAT v35 (44). The best hits with more than 50% of the query coverage were used as the cut-off, considering that approximately 40% of the Lyosin is derived from the highly conserved MYL4. To investigate the intactness of exon L of Lyosin, the sequences of BLAT hits were retrieved with upstream and downstream of 600 nucleotides. The coding sequences from the start to stop codon of more than 200 codons were then extracted using the getorf program in EMBOSS (45). The resulting sequences were aligned and were manually checked to retrieve the intact exon L meeting the following criteria: (1) aligned with the Lyosin sequences previously identified, (2) including the 5’-splice site in frame, and (3) retaining more than 200 codons after trimming of putative introns. The codon alignments of the obtained intact exon L sequences are included in **Dataset S4**. To trace the losses of Lyosin phylogenetically, a taxonomic tree of the species used in the analysis was retrieved from TimeTree 5 on December 7, 2023 (Kumer et al. 2022). The obtained trees were visualized with the annotations of the BLAT result and the intactness of exon L using ggtree v3.10.0 (46).

### Molecular evolutionary analysis

The dN/dS ratio (ω) was estimated using the codeml program in PAML v4.8 (47) based on the codon alignment of the ORF1-like exon L sequences (**Dataset S4**) under the one-ratio ω model: M0. The likelihood ratio tests were conducted by comparing models of ω=1 and estimated ω to test the purifying selection. To detect positive selection, we performed the likelihood ratio tests to compare two pairs of site-specific models (neutral model vs. positive selection model): M1a vs. M2a and M7 vs. M8. The sites under positive selection were identified by the Bayes empirical Bayes procedure on M8.

### Transcriptome analysis

The tissue transcriptomic data of green anoles (SRP102989: from SRR5412144 to SRR5412173) were downloaded (48). The FASTQ files were trimmed using fastp v0.23.2 (49) and mapped to the green anole reference genome (rAnoCar3.1.pri, GCA_035594765.1) using STAR v2.5.2b (50) with “--outSAMattributes NH HI NM MD XS AS --outFilterMultimapNmax 500” options. Genome-guided transcript assemblies using StringTie2 v2.1.6 (51) were conducted for each sample. The generated GTF files were merged using StringTie2 with the “--merge” option. Reads mapped to features in the merged GTF file were counted using featureCounts v2.0.1 (52) with the “-T 8 -s 2 -t exon -g gene_id -J” options. The transcripts of *MYL4* and *Lyosin* were identified by the sequence similarity search. To quantify the splicing, the number of reads spanning exon 2 to 3 (CM069372.1:35022971-35028148) or exon L to 3 (CM069372.1:35026283-35028148) was obtained from the splicing junction tables included in STAR outputs.

### Animals

An adult male Madagascar ground gecko (*Paroedura picta*) was provided by the Animal Resource Development Unit, RIKEN CLST. The experiments were performed in accordance with the regulations for animal experiments approved by the Nagoya University animal experiment committee and the Guidelines for the Proper Conduct of Animal Experiments (Science Council of Japan).

### RT-PCR

Total RNA from the heart and testis of the Madagascar ground gecko was purified using ISOGEN (#311-02501, Nippon Gene) and Direct-zol RNA Miniprep kit (#R2050, Zymo Research). The cDNA was synthesized from total RNA using Verso cDNA synthesis kit (#AB1453A, Thermo Fisher Scientific). PCR was performed with 30 cycles of amplification (10 s at 98°C, 5 s at 60°C, and 5 s at 68°C) by KOD One Master Mix (#KMM-101, TOYOBO). A set of primer 1 (5’-AAGCAGGCTGCCACCATGGCCCCCAAAAAGCCGGA-3’) and primer 2 (5’-ACAAGAAAGCTGGGTTTAAGCGTAATCCGGAACATCGTATGGGTAGCCAGACATGATGTGTTTGAC-3’), and a set of primer 3 (5’-AAGCAGGCTGCCACCATGAAAATGCCAAACAAGTCCAC-3’) and primer 2 were used for amplification of *MYL4* and *Lyosin*, respectively.

### Plasmids

To construct the MYL4 and Lyosin expression plasmids (pPB-MYL4-HA and pPB-Lyosin-HA), the C-terminally HA-tagged MYL4 and Lyosin were amplified from cDNA. The mammalian expression plasmid (#VB900088-2265rnj, VectorBuilder) was linearized by inverse PCR, and the PCR product of MYL4 or Lyosin was inserted into the linearized plasmid. The nucleotide sequences of inserts were determined by Sanger sequencing (Eurofins Genomics). For construction of the reporter plasmid for L1 retrotransposition assay (pL1-spPuro), fragments of human L1 ORF1 and ORF2 from the EF06R plasmid were amplified by PCR and cloned into pcDNA3.1(+) (#V79020, Invitrogen) with the puromycin resistant gene split by an intron. EF06R was a gift from Eline Luning Prak (#42940, Addgene) (53). Sequence data for all plasmids with primers were included in **Dataset S6**. All PCRs described above were carried out using KOD One Master Mix (#KMM-101, TOYOBO), and all fragment ligation reactions were performed using NEBuilder HiFi DNA Assembly Master Mix (#M5520AA, New England Biolabs).

### Western blotting

293T human embryonic kidney cells (#RCB2202; Riken BioResource Research Center) were seeded on 24-well plates. The next day, cells were transfected with pPB-MYL4-HA or pPB-Lyosin-HA using Avalanche Everyday Transfection Reagent (#EZT-EVDY-1; EZ Biosystems, College Park, MD, USA). The cells were lysed in sample buffer for SDS-PAGE (#09499-14, Nacalai Tesque) 24 hours after transfection. SDS/PAGE was performed, and peptides were transferred from the gel to polyvinylidene difluoride membranes. The membranes were reacted with rabbit anti-HA antibody (#361, Medical & Biological Laboratories) and mouse anti-alpha-tubulin antibody (#66031-1-Ig, Proteintech) were used as primary antibodies. Goat anti-rabbit antibody (#32460, Invitogen) and goat anti-mouse IgG antibody (#32430, Invitrogen) were used as second antibodies. Signals were detected using Super Signal West Femto System (#34095, Thermo Fisher Scientific).

### Immunofluorescent assay

One of HA-tagged protein expression plasmids (pPB-MYL4-HA, pPB-Lyosin-HA, or pPB-L1ORF1p-HA) and pPB-L1ORF1p-FLAG were transfected into 293T cells seeded into a slide chamber (#192-008, Watson) using Avalanche Everyday Transfection Reagent. At 24 hours post transfection, the cells were fixed with 4% paraformaldehyde phosphate buffer for 15 min at room temperature. The cells were incubated with blocking buffer [PBS, 5% (w/v) BSA, 0.5% (v/v) Triton X-100, 0.1% (v/v) Tween 20] for 15 min at room temperature. The cells were reacted with rabbit anti-HA antibody and mouse anti-FLAG antibody (#F3165, Sigma-Aldrich) diluted with antibody reaction buffer [PBS, 1% (w/v) BSA, 0.1% (v/v) Tween 20] for one hour at room temperature. After washing with PBS, the cells were incubated with goat anti-rabbit IgG with Alexa Fluor 594 (#A11012, Invitrogen) and goat anti-mouse IgG with Alexa Fluor 488 (#A32723, Invitrogen) diluted with the antibody reaction buffer for one hour at room temperature. After washing with PBS, the cells were observed with fluorescence microscopy (#BZ-X810, KEYENCE).

### L1 retrotranspositon assay

293T cells were seeded on a 24-well plate, and the next day, the reporter plasmid, pL1-spPuro, was transfected into cells with one of the protein expression plasmids (pPB-EGFP, pPB-MYL4-HA, pPB-Lyosin-HA, or pPB-MOV10) using Avalanche Everyday Transfection Reagent. After 72 hours of transfection, cells were passaged to a 6-well plate with 1 µg/ml of puromycin. After 14 days, the cells were fixed with 4% paraformaldehyde phosphate buffer and stained with 1% Crystal Violet.

### Screening vertebrate RefSeq proteins for ORF1p chimeric isoforms

To identify vertebrate genes with alternative splicing isoforms containing ORF1p-like domains, vertebrate RefSeq proteins (release 220) were downloaded from NCBI on October 12, 2023. Proteins labeled “isoform” were retrieved and were then subjected to a sequence homology search using MMseqs2 (54) against 147 LINE ORF1 proteins of Dfam3.6 (55). The E-value cutoff was set at 1E-10, and the alignment coverage to ORF1p was required to be more than 50%. Then, protein-coding genes with both ORF1p-like isoforms and non-ORF1p-like isoforms (i.e., isoforms that are not similar to ORF1p) were collected from each species. To identify evolutionarily conserved chimeric isoforms, we retrieved these protein-coding genes that were commonly identified in two or more genera. At this point, we identified three protein genes (*MYL4, RFX5, UNC13C*) from non-mammalian RefSeq proteins and three protein genes (*NUP42, USP4, ERVFC1*) from mammalian RefSeq proteins (**Dataset S7**). Of these, *ERVFC1* was identified in marmoset (*Callithrix jacchus*) and vampire bat (*Desmodus rotundus*); however, those evolutionary relationships were not considered to be orthologous based on their flanking genes. Presumably the different L1 ORF1 loci were annotated as genes with the same name. Next, we used the NCBI gene browser (https://www.ncbi.nlm.nih.gov/gene/) to investigate whether the ORF1p-like isoforms were derived from an alternative exon (**Fig. S5**). Unexpectedly, the ORF1p-like exon in *UNC13C* was a canonical coding exon that is also found in human and mouse. In three species, *UNC13C* contained the isoforms lacking this ORF1p-like exon and was identified as a gene with an alternative ORF1p-like isoform in our workflow (**Fig. S6A**). Thus, *UNC13C* is likely to have a canonical coding exon homologous to L1 ORF1 (**Fig. S6B**). This is interesting; however, *UNC13C* was outside of the scope of this study and no further analysis was performed. As a result, four genes, *MYL4, RFX5, NUP42*, and *USP4*, were identified as genes having an alternative ORF1p-like isoform (**Fig. 5**).

## Acknowledgments

We are grateful to Dr. Hiroshi Kiyonari (RIKEN CLST, Japan) for kindly providing of an adult male Madagascar ground gecko (*Paroedura picta*). This work was supported by Grant-in-Aid for JSPS fellows JP23KJ1055 to K.K. and JSPS KAKENHI JP20K06775 to S.N. The supercomputing resource was partially supported by the NIG supercomputer at ROIS National Institute of Genetics.

## Author contributions

K.K. and S.N. designed research; K.K. performed research; K.K., I.K., and S.N. analyzed data; K.K., K.I., and S.N. wrote paper.

## Data Availability

All data generated during this study are included in the main text and supplementary datasets. The codes used in this study are available at (https://github.com/KoichiKitao/LINE_fusion_2024).

---

**Fig. S1.**
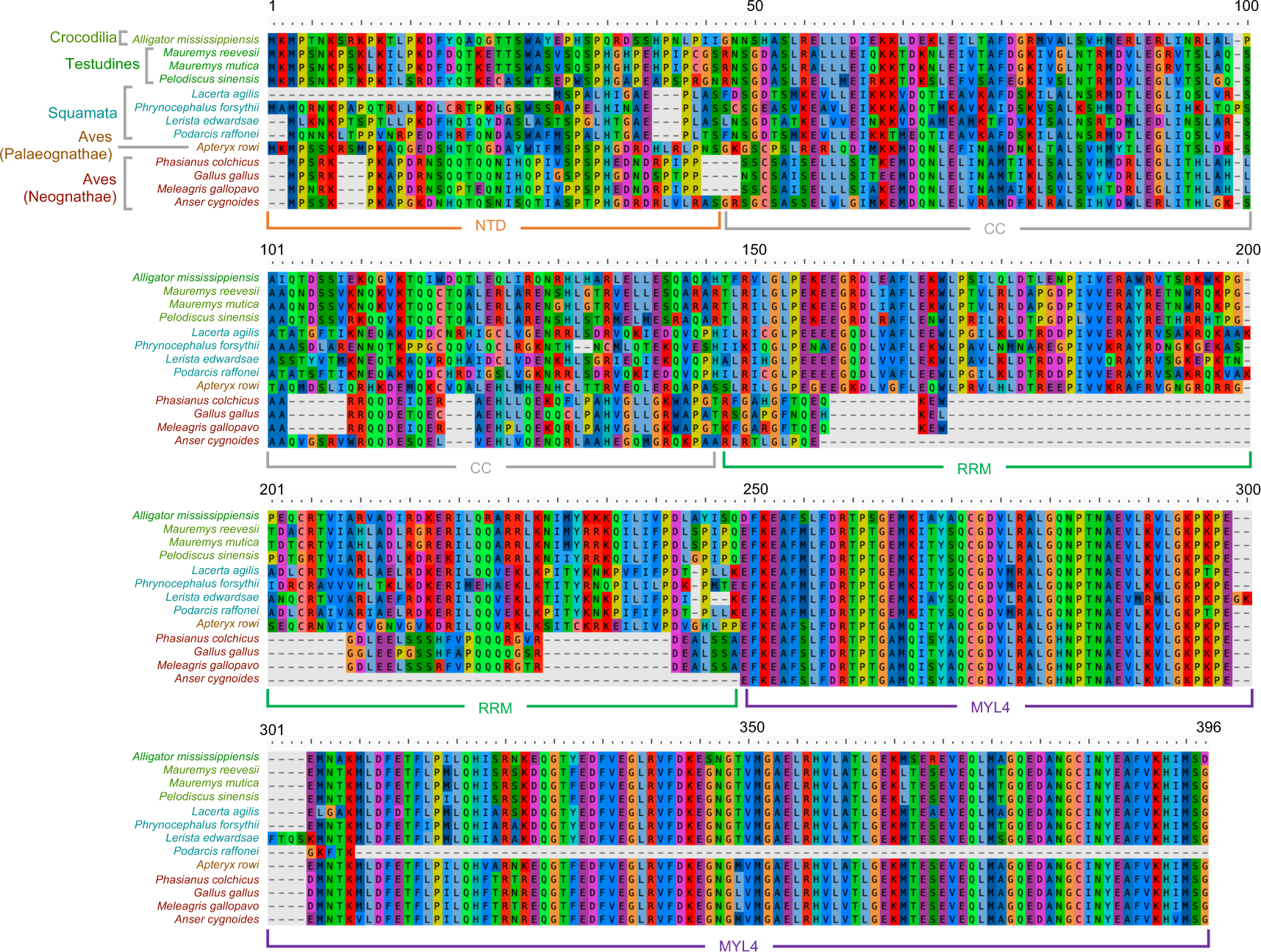
An alignment of Lyosin-like proteins obtained from NCBI BLASTP search. The accession numbers of the proteins were listed in **Dataset S1**.

**Fig. S2.**
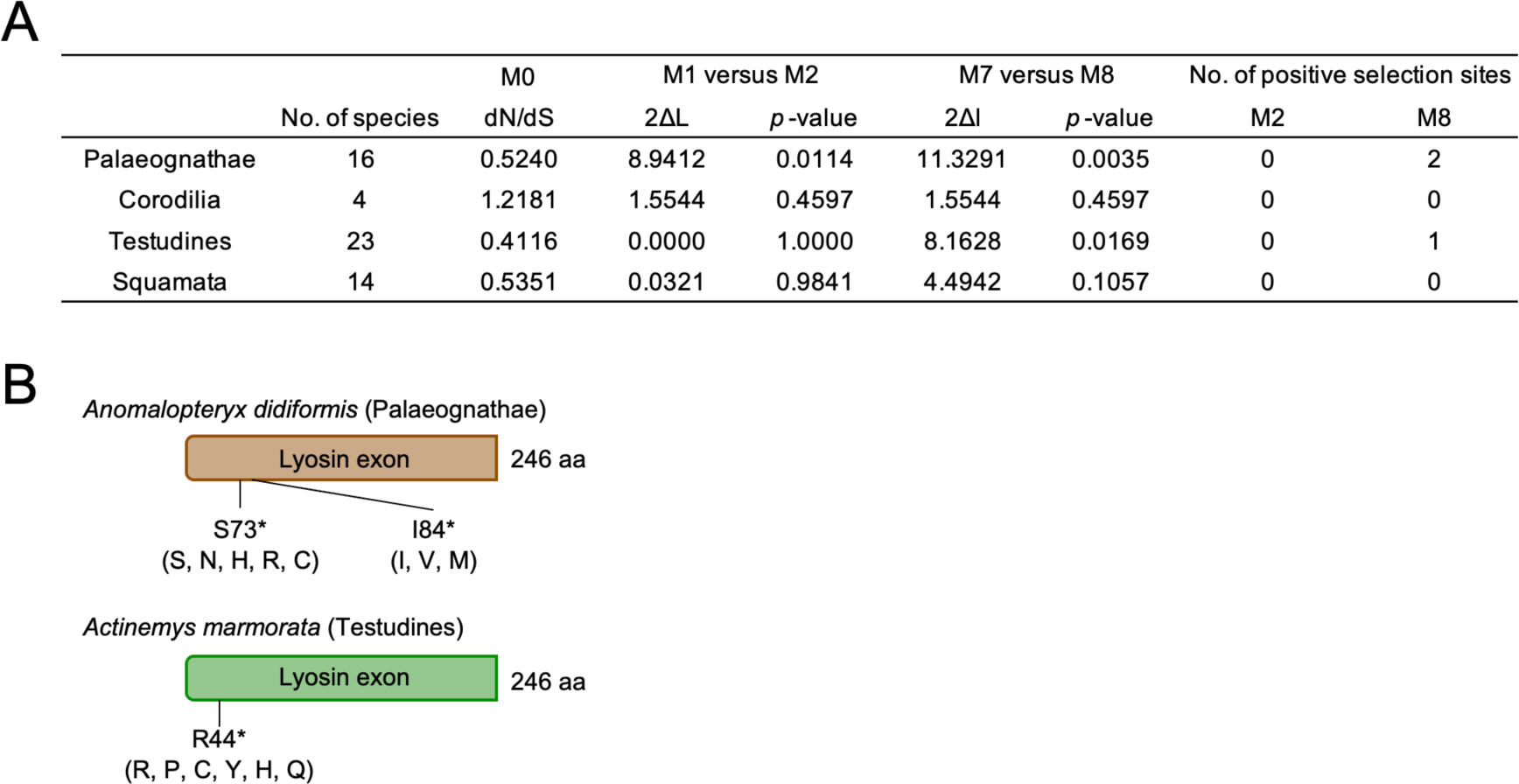
Analysis of the nonsynonymous and synonymous codon substitution (dN/dS) ratio. (A) 2ΔL indicates a two-fold difference in the natural log values of the maximum likelihood ratio from pairwise comparisons of the different models. The *p*-value indicates the confidence in rejecting the natural models (M1a or M7) in favor of the positive selection model (M2a or M8) using Pearson’s chi-squared test. Codons under positive selection were identified with a posterior probability of 95% by Bayes empirical Bayes (BEB) analysis in M8. (B) The sites under positive selection were illustrated based on the Lyosin exon L of representative species. Amino acids other than the representative sequences were shown in brackets.

**Fig. S3.**
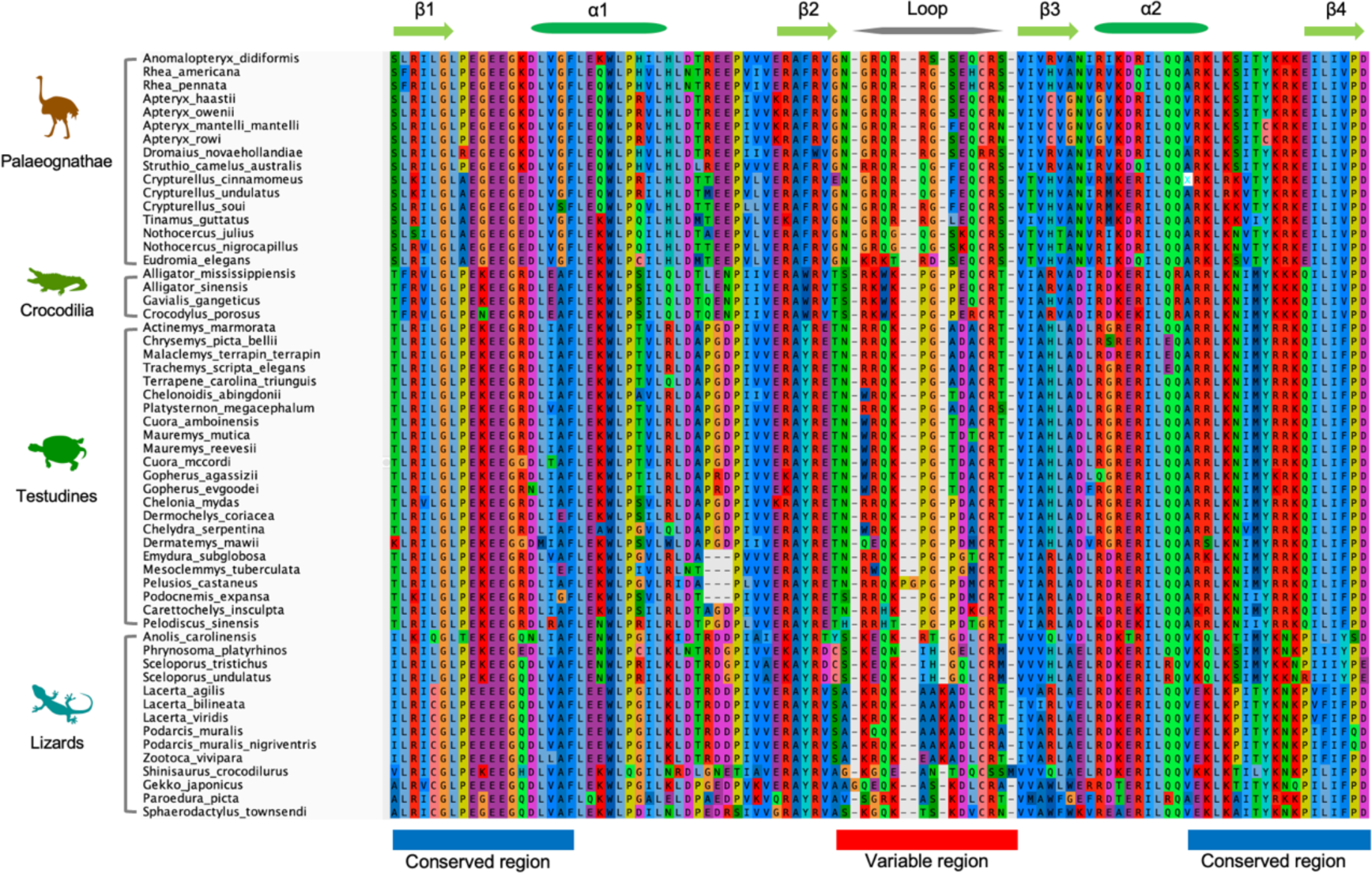
An alignment of the RRM domains of Lyosin. The upper part of the alignment shows the structure of the RRM domain (see Fig 1D).

**Fig. S4.**
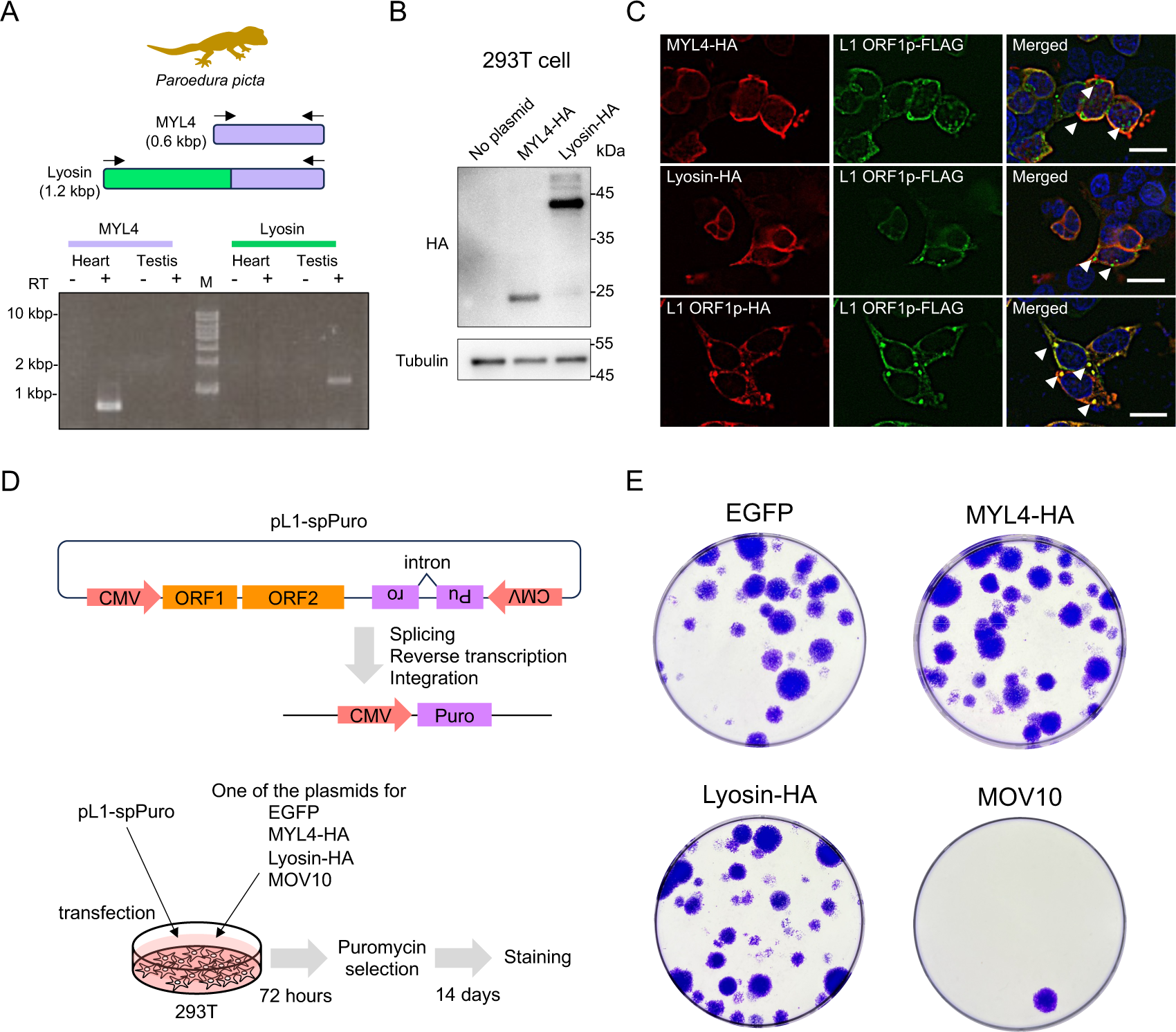
Molecular characterization of the Lyosin protein. (A) RT-PCR targeting *MYL4* and *Lyosin*. RNA was extracted from the heart and testis of a sexually matured male Madagascar ground gecko. Samples without reverse transcriptase (RT-) were analyzed as negative controls. (B) Western blotting of MYL4-HA and Lyosin-HA expressed in human 293T cells. (C) Fluorescence immunostaining images in human 293T cells. White arrows indicate L1 ORF1p puncta. The scale bars represent 20 µm. (D) Illustration of the mechanism of L1 retrotransposition assay. (E) Results of L1 retrotransposition assay. The indicated genes and L1 reporter were co-expressed in 293T cells. L1 retrotransposition results in cell colonies resistant to puromycin. MOV10 is used as a positive control that was reported to suppress L1.

**Fig. S5.**
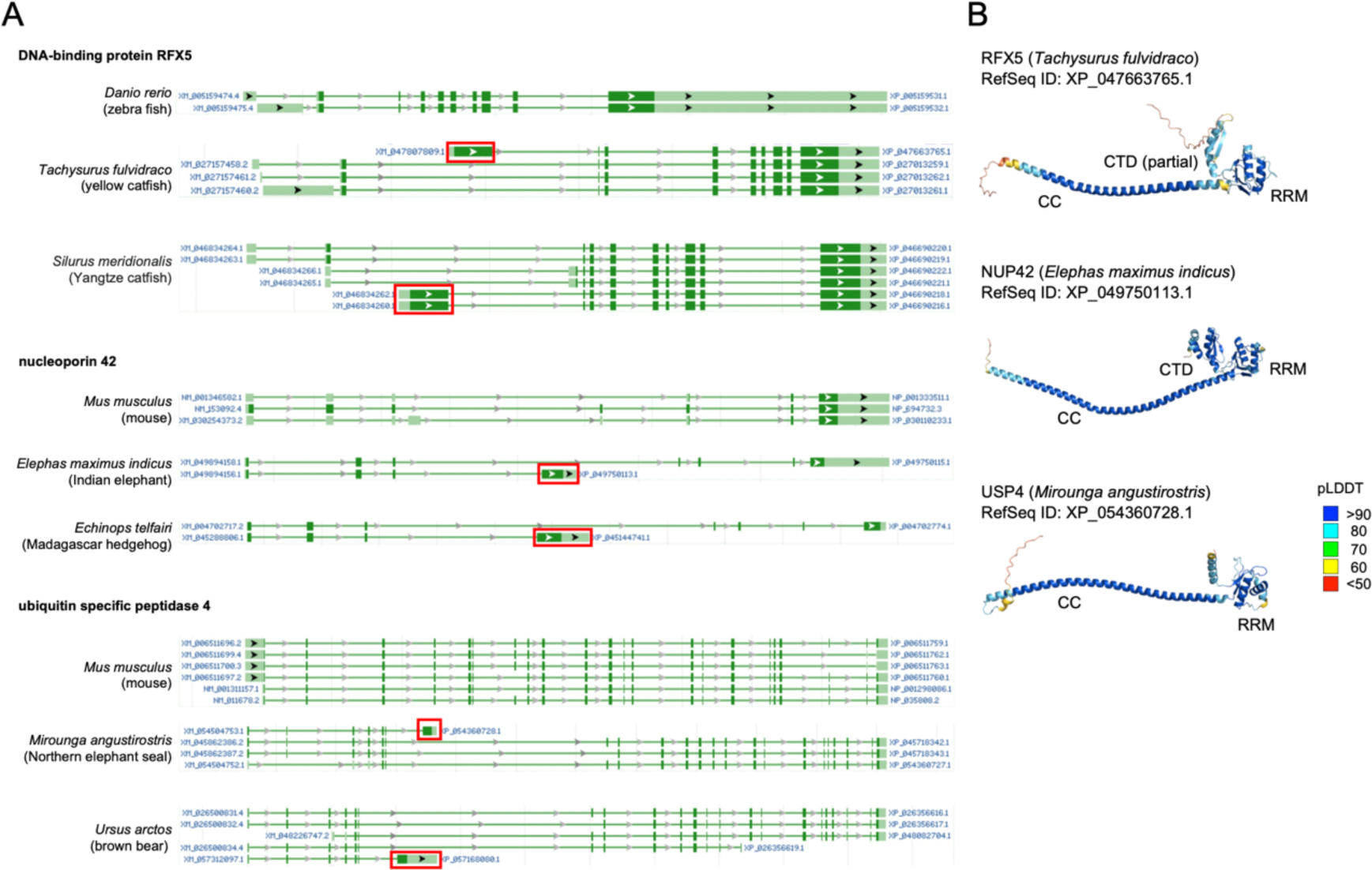
Other genes with ORF1p-like isoforms. (A) Screenshots from the NCBI gene browser. The squared exons are non-canonical alternative exons encoding ORF1p-like proteins. (B) Protein structures of the ORF1p-like amino acid sequence encoded by the non-canonical exons were predicted using AlphaFold2. Predicted domains specific to L1 ORF1p were noted: coiled-coil (CC), RNA recognition motif (RRM), and C-terminal domain (CTD).

**Fig. S6.**
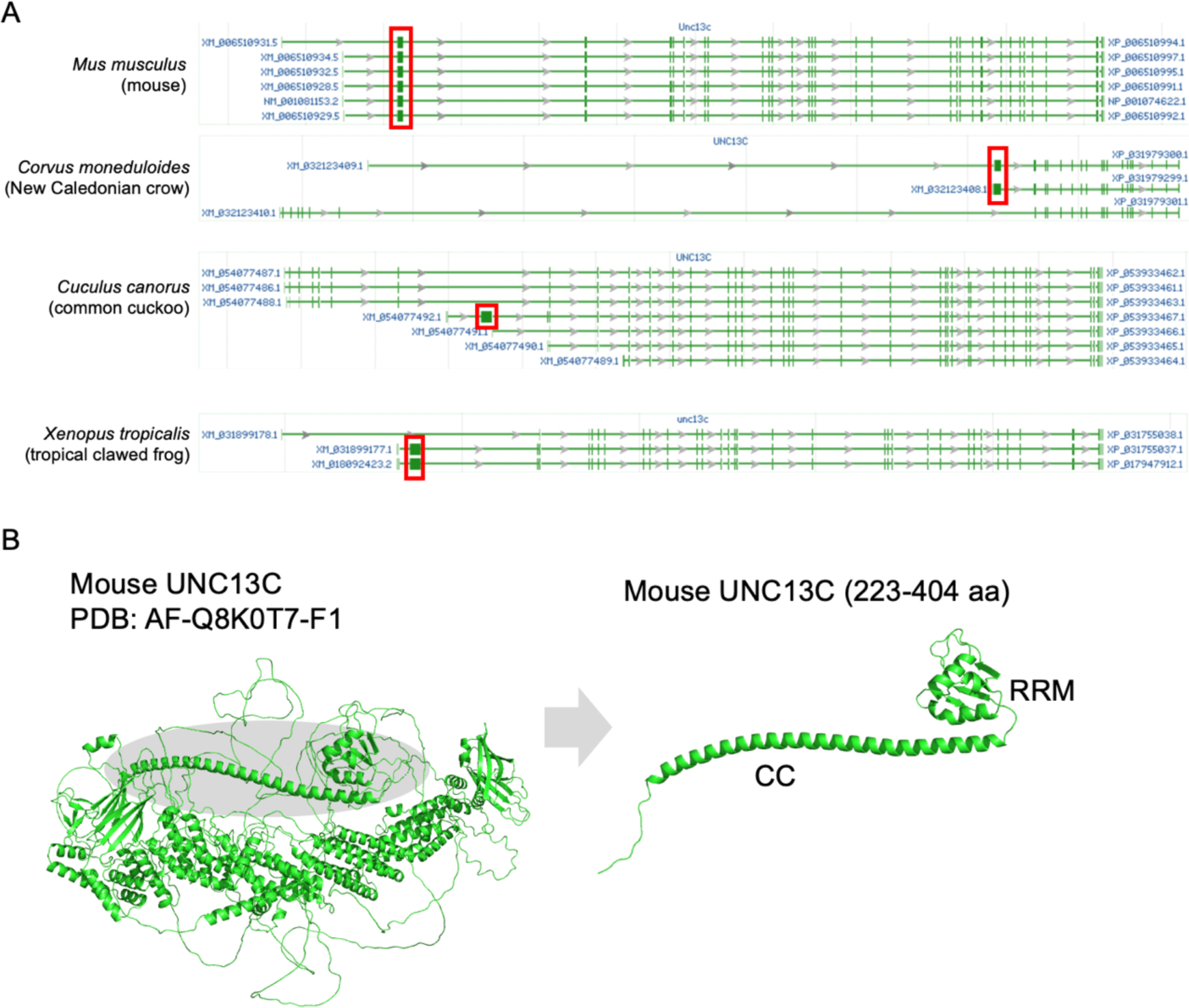
A canonical exon of *UNC13C* encodes a protein similar to L1 ORF1p. (A) Exons encoding ORF1p-like amino acids are enclosed in squares. In mouse, all splicing variants of *Unc13c* contain the exon encoding ORF1p-like amino acids. In the three species below, splicing variants without ORF1p-like exons are annotated. (B) The 3D structure of mouse UNC13C predicted by Alphafold2 (PDB: AF-Q8K0T7-F1). On the left is the predicted structure of full-length UNC13C, while on the right is the partial structure corresponding to the L1 ORF1p-like region. CC, coiled-coil; RRM, RNA recognition motif.

